# SpatialArtifacts: a computational framework for tissue artifact detection in spatial transcriptomics data

**DOI:** 10.64898/2026.05.15.725260

**Authors:** Jiali Harriet He, Jacqueline R. Thompson, Michael Totty, Stephanie C. Hicks

## Abstract

Spatial transcriptomics data are frequently compromised by technical artifacts, such as dry patches, tissue lifting, and uneven reagent coverage, which manifests as regions with low UMI counts, in particular at tissue borders. It can often be challenging to identify these regions using existing quality control methods. Here, we present `SpatialArtifacts`, a framework that combines median absolute deviation (MAD)-based outlier detection with mathematical morphology operations to identify and classify spatially contiguous tissue artifacts. Focal operations including 3×3 fill, 5×5 outline, and star-pattern connectivity link low-quality spots while preserving true biological domains. We use a hierarchical classification system to distinguish edge versus interior artifacts and large versus small regions, enabling downstream removal or targeted manual review. We demonstrate the performance of our method in human hippocampus, dorsolateral prefrontal cortex, and colorectal cancer tissues using 10x Genomics Visium and VisiumHD platforms. Our `SpatialArtifacts` package is freely available on Bioconductor at https://bioconductor.org/packages/SpatialArtifacts and on PyPI at https://pypi.org/project/spatial-artifacts/.

## 1 Introduction

Spatial transcriptomics has transformed biological research by enabling the study of gene expression in its native tissue context [1]. However, the utility of these datasets is frequently affected by technical artifacts that arise during sample preparation, which can compromise downstream analyses [2]. A common source of these artifacts stems from the physical distortion of the tissue. When tissue lifts or detaches from the slide surface, reagents might distribute unevenly leading to a reduction in the amount of data captured on the edges. Similarly, tissue folding during sectioning or inadequate drying can produce inconsistent loss of captured gene expression. Regardless of the underlying cause, these artifacts often share a common spatial signature: geographically continuous regions of the tissue with abnormally low gene expression counts or elevated mitochondrial expression. Ignoring these artifacts can lead to a misclassification of biologically distinct niches in unsupervised clustering, differential expression, and cell-cell interactions [3, 4].

Existing computational quality control (QC) methods for spatial transcriptomics have been developed to handle some of these artifacts. They typically use global threshold-based filtering that evaluates the spot in isolation relative to all other spots. In constrast, spatially-aware quality control tools such as SpotSweeper [5] have been developed to evaluate the quality of the spot (or cell) in the context of its closest set of nearest neighbors (NN), for example *k*=6 NN, making it more robust to biological heterogenity of the tissue. SpotSweeper also provides artifact categorization for specific artifact types, such as “hangnails”. Another tool, BLADE (Border, Location, and edge Artifact DEtection) [6] takes a complementary approach by applying a *t*-test to determine whether edge regions are statistically distinct from the interior tissue and removing a fixed-depth ring of spots when a significant difference is detected. Importantly, there are different types of signal loss in spatial transcriptomics, such as (i) low expression due to biases such as PCR amplification bias associated with the synthetic DNA barcodes, or (ii) tissue artifacts caused by physical damage. While the former tends to be isolated, the latter is often geographically coherent and whose extent is defined by the physical processes that caused them, such as surface tension, reagent flow, and tissue adhesion, rather than by any predefined geometric rule. Despite these advances, important gaps remain. SpotSweeper’s local quality control test statistics were not designed to identify irregularly bounded regions, which are often the shapes of tissue artifacts. BLADE, while effective at edge correction, operates at the level of entire margins, rather than at in individual spots, leading to unnecessary removal of high-quality tissue.

Here, we present SpatialArtifacts, a package implemented in both R (Bioconductor) and Python (PyPI) that addresses these limitations through spatial pattern recognition (**Figure 1**). Identifying the spatial footprint of these tissue artifacts requires an approach that can trace irregular boundaries across the tissue section. We reasoned that morphological image processing [7], methods with deep roots in computer vision and medical analysis, would be ideal for this task. Mathematical morphological operations such as dilation, erosion, gap filling, and connected-component labeling were developed precisely to segment objects with sharp, irregular boundaries, properties shared by pathological lesions in histopathology and MRI images, similar to the artifact regions we seek to identify here. Our method first identifies “seed” low-quality spots using median absolute deviations (MADs) applied at the level of a tissue section, then employs a series of morphological operations to expand these seeds into coherent patches that reflect the irregular geometry of technical artifacts. This approach is artifact-agnostic and can delineate spatially contiguous low-quality regions using mathematical morphology operations, regardless of its shape or origin. By enabling precise, spot-level artifact boundaries, SpatialArtifacts maximizes the retention of high-quality data while minimizing false removals, improving the reliability of spatial transcriptomics datasets.

**Figure 1:**
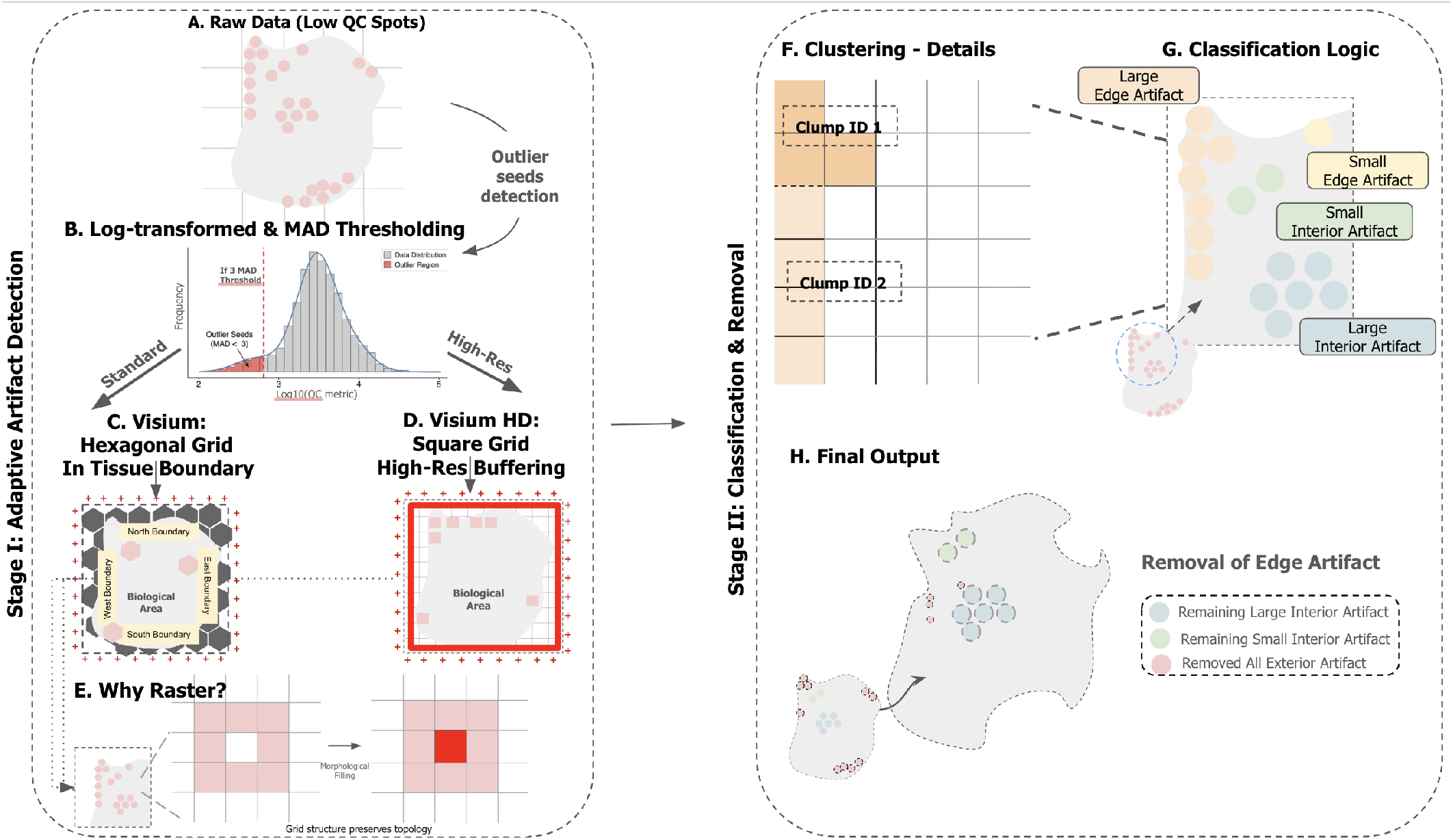
Overview of SpatialArtifacts for mathematical morphology-based tissue artifact detection. The SpatialArtifacts pipeline consists of two stages. In **Stage I (Adaptive Artifact Detection)**, (**A**) Outlier spots or “seeds” (pink) in spatial transcriptomics data are identified using median absolute deviations (MADs). (**B**) Distribution of QC metric (log_10_ scale) with a threshold (dashed line) classifying outlier seeds from normal tissue. Platform-specific detection strategies are then applied. (**C**) Standard Visium: hexagonal grid with biological tissue boundary (`in_tissue` flag) used for edge detection. Morphological image processing connects spatially adjacent outliers. (**D**) VisiumHD: square grid with physical boundary buffering (red frame, default 80*µ*m) used for high-resolution edge detection, adapting to irregular tissue contours. (**E**) Rasterization converts irregular spot coordinates into unified grid space, preserving spatial topology for morphological image processing (see Figure S1 for kernel details). In **Stage II (Classification and Removal)**, (**F**) Graph-based clustering identifies connected outlier components (Clump ID 1, Clump ID 2, etc.) after morphological aggregation. (**G**) Hierarchical classification assigns artifacts to four categories based on location (edge versus interior) and size (large versus small): (i) large edge, (ii) small edge, (iii) large interior, and (iv) small interior. (**H**) Final predicted tissue artifacts.

## 2 Results

### 2.1 A mathematical morphology-based framework for identifying spatial artifacts

SpatialArtifacts implements this idea in a two-stage pipeline (**Figure 1**). In Stage I (Adaptive Artifact Detection), we use MAD-based thresholding applied to log-transformed QC metrics across spots at the tissue section level (**Figure 1A-B**). These outlier spots serve as initial “seeds” of potential artifact regions. Because the raw per-spot QC metrics often produce a fragmented picture of spatially continuous damage, the algorithm converts the platform-specific spatial coordinates, such as hexagonal spots or square bins (**Figure 1C-D**), into a unified raster grid, transforming the problem into one observed in image processing. Next, a series of morphological focal operations, including 3×3 gap filling, 5×5 outline detection, and star-pattern connectivity, are applied to connect neighboring outlier seeds into coherent patches that reconstruct the true spatial geometry of the artifact, while preserving genuine biological boundaries (**Figure 1E, Figure S1**).

In Stage II (Classification and Removal), we use graph-based clustering to connect outliers into discrete tissue artifact regions (**Figure 1F**). Then, these regions are classified hierarchically considering two dimensions, namely the spatial location relative to the tissue boundary (*edge* versus *interior*) and the cluster size (*large* versus *small*) yielding four categories: (i) large edge, (ii) small edge, (iii) large interior, and (iv) small interior (**Figure 1G-H**). This classification reflects both the likely etiology and the confidence of removal: edge artifacts are the most common consequence of physical damage and can be easily removed, while large interior artifacts are flagged for manual review given their potential overlap with biologically meaningful low-expression regions. The result is a precise, spot-level artifact delineation that maximizes retention of high-quality tissue while minimizing false removals (**Figure 1H**).

### 2.2 SpatialArtifacts identifies tissue artifacts in human hippocampus tissue

The human hippocampus has a complex cytoarchitecture and broad dynamic range of transcriptional activity. We applied SpatialArtifacts to a postmoretum human hippocampus tissue section profiled on the 10x Genomics Visium platform [3, 8] (**Figure 2**). Total unique molecular identifiers (UMIs) varied substantially across the tissue (**Figure 2A**), reflecting biological heterogeneity and technical noise. Using a fixed QC threshold (UMI *<* 1000) [5] as an illustrative upper bound, we found low-quality spots scattered throughout the tissue, including regions where lower UMI counts are expected due to differences in cell composition throughout the hippocampus (**Figure 2B**). At the same time, using three commonly-used fixed QC thresholds (UMI *<* 300, UMI *<* 500, and UMI *<* 1000) as comparison, we found that lower thresholds such as UMI *<* 300 failed to detect the majority of artifact regions, while UMI *<* 500 most closely approximated SpatialArtifacts results for this sample but would not generalize across samples with different expression profiles. UMI *<* 1000 flagged the most spots but also included extensive biologically meaningful low-expression regions (**Figure S2G–I**). This demonstrates that no single global threshold can reliably distinguish technical artifacts from biologically meaningful low-expression regions, motivating the need for a spatially-aware approach. And so, for consistency, we use UMI *<* 1000 as the global threshold comparison for standard Visium samples throughout this manuscript and UMI *<* 500 for VisiumHD samples, reflecting the lower sequencing depth per bin at higher spatial resolution.

**Figure 2:**
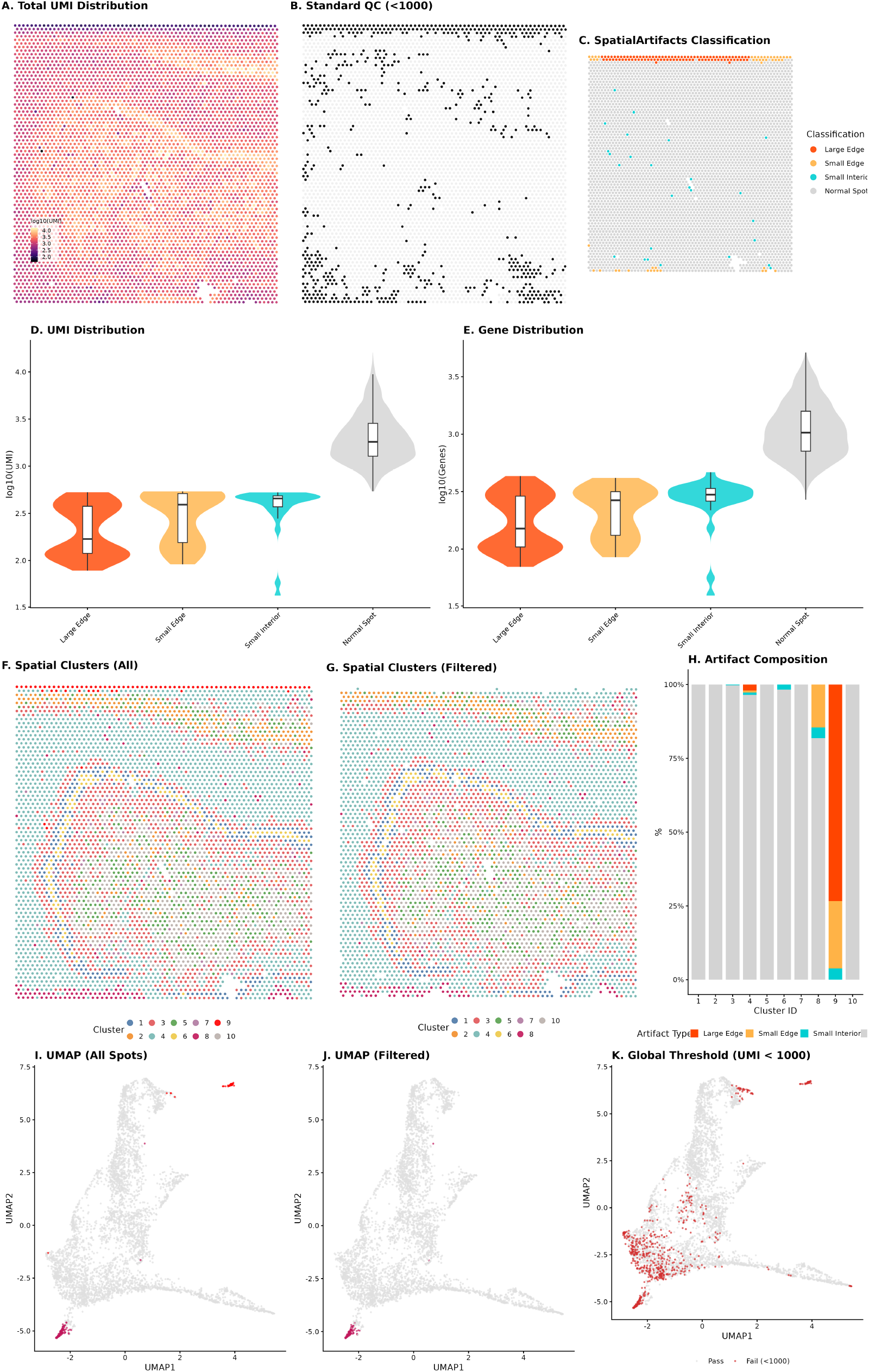
SpatialArtifacts identifies tissue artifacts in human hippocampus tissue. Tissue sample `V11L05-335_C1` from a public dataset [8]. (**A**) Heat map of total UMI (log_10_ scale). (**B**) Spots flagged as low-quality (black) using standard global QC thresholds (UMI *<* 1000). (**C**) SpatialArtifacts classification identifies spatially coherent artifacts at tissue boundaries and folds: Large Edge (orange), Small Edge (yellow), Small Interior (cyan), with Normal Spots (gray) preserved. (**D–E**) Violin plots for the distribution of UMI counts (D) and number of detected genes (E) across artifact and non-artifact spots. Large Edge artifacts show reduced expression relative to Normal Spots (median log_10_(UMI) ≈ 2.2 vs 3.2). (**F–G**) Spatial clustering before (F) and after (G) artifact removal. Raw data contains 10 clusters; filtered data retains 8 biological clusters with improved spatial coherence. Artifact-dominated Cluster 9 (red) is entirely eliminated after filtering, and partially affected Cluster 8 (pink) retains only non-artifact spots. (**H**) Artifact composition analysis quantifies contamination per cluster. Cluster 9 consists entirely of predicted artifact spots (100%), while Cluster 8 contains 18.2% artifact spots. (**I–J**) UMAPs before (I) and after (J) filtering. Artifact-driven clusters (red/pink points) form distinct non-biological manifolds in raw data; these are eliminated after targeted removal, restoring topological coherence. (**K**) Global thresholding comparison (UMI *<* 1000, red points) removes substantially more spots.

In contrast, SpatialArtifacts identified spots concentrated mainly at the edges of the tissue, preserving interior regions with naturally lower expression (**Figure 2C**). To confirm that the detected artifacts represent technical rather than biological signal loss, we examined the spatial expression of established hippocampal marker genes (SNAP25 and GFAP). Edge artifact spots showed complete loss of marker gene expression (median log-normalized expression = 0 for both SNAP25 and GFAP), whereas Normal Spots retained robust expression (median = 2.48 for both markers). Interior artifact spots showed intermediate signal loss (median GFAP = 1.23), consistent with partial technical degradation rather than biologically meaningful low expression (**Figure S2D–F**). Although all artifact spots exhibited low UMI counts, the key distinction from standard global thresholding lies in spatial specificity: of the 659 spots flagged by a UMI *<* 1000 threshold, 492 had a median UMI of 840 and were correctly preserved by SpatialArtifacts as biologically meaningful low-expression regions (**Figure S2C**). Performing spatial clustering revealed that spots identified with SpatialArtifacts were mostly from one or two clusters, with cluster 9 consisting entirely of artifact spots (100%) and cluster 8 containing 18.2% artifact spots (**Figure 2H**).

After filtering out all spots flagged by SpatialArtifacts (**Figure 2G**), the UMAP was better integrated (**Figure 2J**). To evaluate the impact of artifact removal on transcriptional variance, we applied spotlevel PCA and used `plotExplanatoryPCs` to assess the contribution of metadata variables to the top 10 principal components. Prior to filtering, artifact classification explained a substantial proportion of variance in PC1 and PC2. After removing artifact spots, the contribution of QC metrics to mid-range PCs (PC4–6) decreased by approximately 100-fold (from ∼0.1% to ∼0.001%), suggesting that these components previously captured artifact-driven technical variation rather than biological signal (**Figure S2A–B**). By comparison, fixed global thresholds removed spots from tissue artifacts and meaningful biological domains indiscriminately (**Figure 2K**).

### 2.3 Enhanced recovery of cortical layer architecture in human DLPFC

The human dorsolateral prefrontal cortex (DLPFC) presents a unique challenge for tissue artifact detection due to the presence of myelinated axons in white matter, which produce naturally low transcript counts [10] (**Figure 3A**). We evaluated SpatialArtifacts using a postmortem human DLPFC sample [9], which includes computational spatial domain labels of six cortical layers plus white matter [10]. This served as a ground truth to determine if artifact removal improved the identification of anatomical domains. Notably, the dataset included 87 spots that the original authors could not assign to any cortical layer (labeled as ‘Unannotated’), presumably due to low quality spot. Using a standard QC threshold (UMI < 1000) (see **Figure S2G–I** for threshold comparison), low-quality spots were scattered throughout the tissue, including in white matter regions (**Figure 3B**).

**Figure 3:**
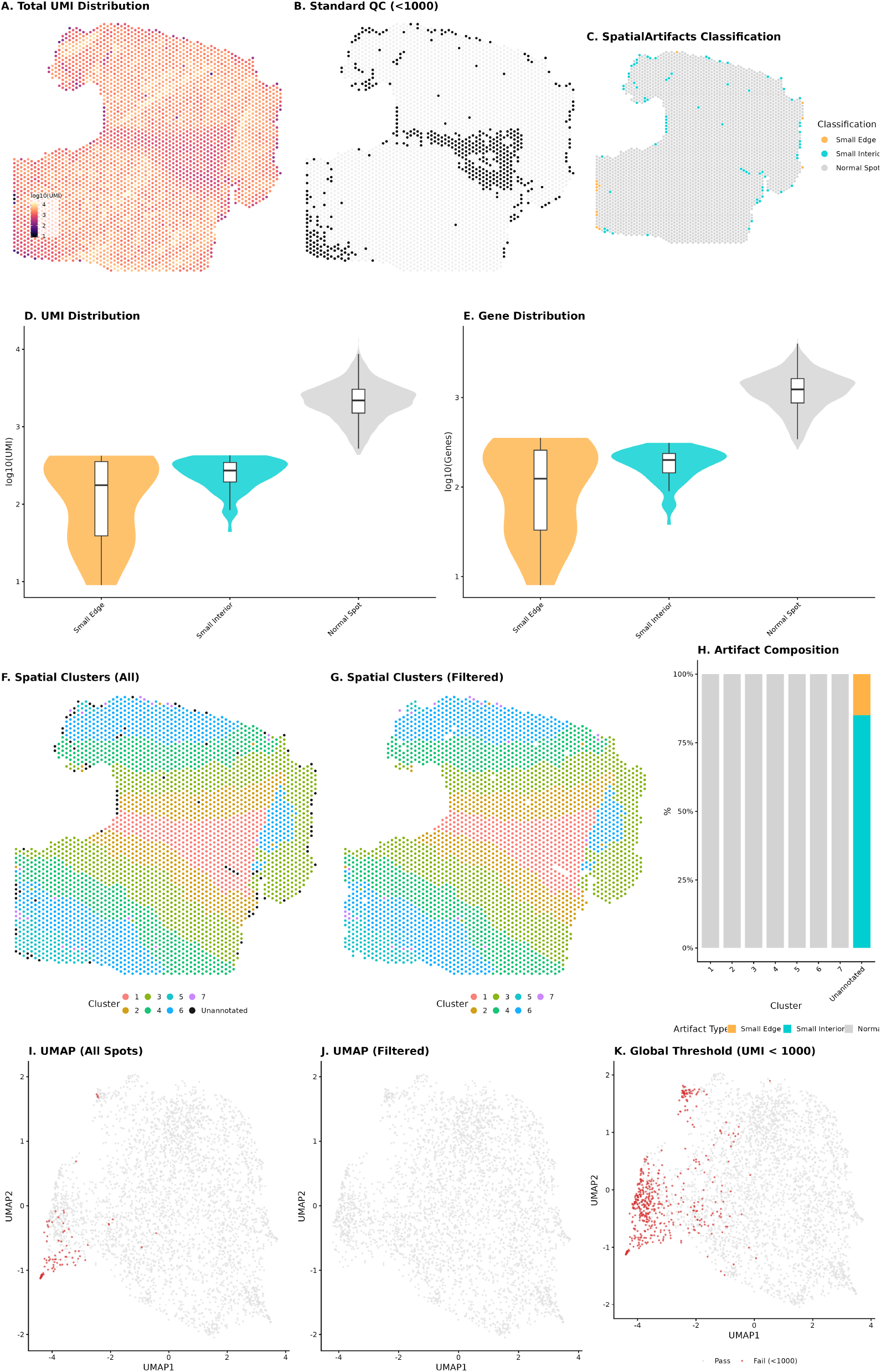
Enhanced recovery of cortical layer architecture in the human DLPFC. Tissue sample `Br8325_ant` from public dataset [9]. (**A**) Heat map of total UMI (log_10_ scale). (**B**) Spots flagged as lowquality (black) using standard global QC thresholds (UMI *<* 1000). (**C**) SpatialArtifacts classification localizes artifacts to tissue edges and fold regions: Small Edge (orange) and Small Interior (cyan), while preserving white matter structural integrity (gray). (**D–E**) Violin plots for the distribution of UMI counts (D) and number of detected genes (E) across artifact and non-artifact spots. (**F–G**) Spatial clustering before (F) and after (G) artifact removal. The 87 ‘Unannotated’ spots (black) are entirely identified as artifacts by SpatialArtifacts and are removed after filtering. (**H**) Artifact composition analysis confirms that the ‘Unannotated’ cluster consists entirely of classified artifacts (100% Small Edge or Small Interior). (**I–J**) UMAP before (I) and after (J) filtering. Artifact-associated spots (red) form a distinct region in raw data that is eliminated after targeted removal. (**K**) Global thresholding (UMI *<* 1000, red) removes extensive regions including substantial portions of valid white matter.

In contrast, SpatialArtifacts (using a MAD > 3) localized tissue artifacts to mostly tissue edges and tissue fold regions (**Figure 3C**), while retaining white matter regions. Compositional analysis revealed a striking concordance between SpatialArtifacts and the expert-curated quality control used in the original study: the 87 spots that remained “Unannotated” by the authors were found to consist entirely of classified artifacts (13 Small Edge and 74 Small Interior spots) (**Figure 3D-G**). This demonstrates that SpatialArtifacts can automate the identification of low-quality regions that otherwise require manual, time-consuming exclusion by domain experts. Also artifacts at tissue folds are called Small Interior instead of Small Edge. This is because the edge identification method finds clusters that touch the slide capture area’s border instead than the tissue boundary itself. Although all artifact spots exhibited low UMI counts, the key distinction from standard global thresholding lies in spatial specificity: of the 476 spots flagged by a UMI *<* 1000 threshold, 389 had a median UMI of 773 and were correctly preserved by SpatialArtifacts as biologically meaningful low-expression regions (**Figure S3C**). The QC metric distributions confirmed that detected artifacts exhibited lower UMI counts and gene detection compared to Normal Spots (**Figure 3D–E**). Examination of cortical marker gene expression revealed that artifact spots retained partial SNAP25 expression (median = 2.84 for Edge, 2.78 for Interior artifacts vs 3.14 for Normal Spots), while GFAP expression was absent in artifact spots (median = 0) compared to Normal Spots (median = 1.42), consistent with selective technical degradation rather than complete signal loss (**Figure S3D–F**).

After filtering out all spots flagged by SpatialArtifacts (**Figure 3G**), the UMAP showed improved integration (**Figure 3J**). Spot-level PCA confirmed that artifact removal reduced the contribution of artifact classification to PC1 and PC2, while the variance explained by PC2 and PC3 increased substantially after filtering (from ∼0.1% to ∼1% and from ∼0.001% to ∼1–10%, respectively), suggesting that biological signal previously masked by technical artifacts became more apparent (**Figure S3A–B**). By comparison, fixed global thresholds removed spots from tissue artifacts and meaningful biological domains indiscriminately, including substantial portions of the white matter (**Figure 3K**).

### 2.4 Computational scalability across platforms and tissue architectures

To demonstrate generalizability across platforms and tissue architectures, we applied SpatialArtifacts to a human colorectal cancer VisiumHD sample [4] at 16 *µ*m resolution following existing workflows for VisiumHD [11]. A map of UMI distribution (**Figure 4A**) revealed complex mucosal architecture with extreme spatial heterogeneity. Standard global thresholding (UMI *<* 500) (see **Figure S2G–I** for threshold comparison) [5] flagged extensive regions across the entire section of tissue, including biologically relevant mucosal crypts and muscularis (**Figure 4B**). In contrast, SpatialArtifacts identified a narrow set of edge artifacts, interior damage sites, and scattered folds as contiguous patches with distinct borders despite the sevenfold increase in spatial resolution relative to standard Visium (**Figure 4C**). The distribution of total UMIs and detected genes per bin was substantially lower in artifact bins compared to normal bins (median UMI = 33–42 across artifact categories vs 2,264 for Normal Spots) (**Figure 4D–E**). To confirm that the detected artifacts represent technical rather than biological signal loss, we examined the spatial expression of colorectal tissue marker genes EPCAM, ACTA2, and CDH1, which showed preserved expression in interior regions correctly retained by SpatialArtifacts (**Figure S4F–H**).

**Figure 4:**
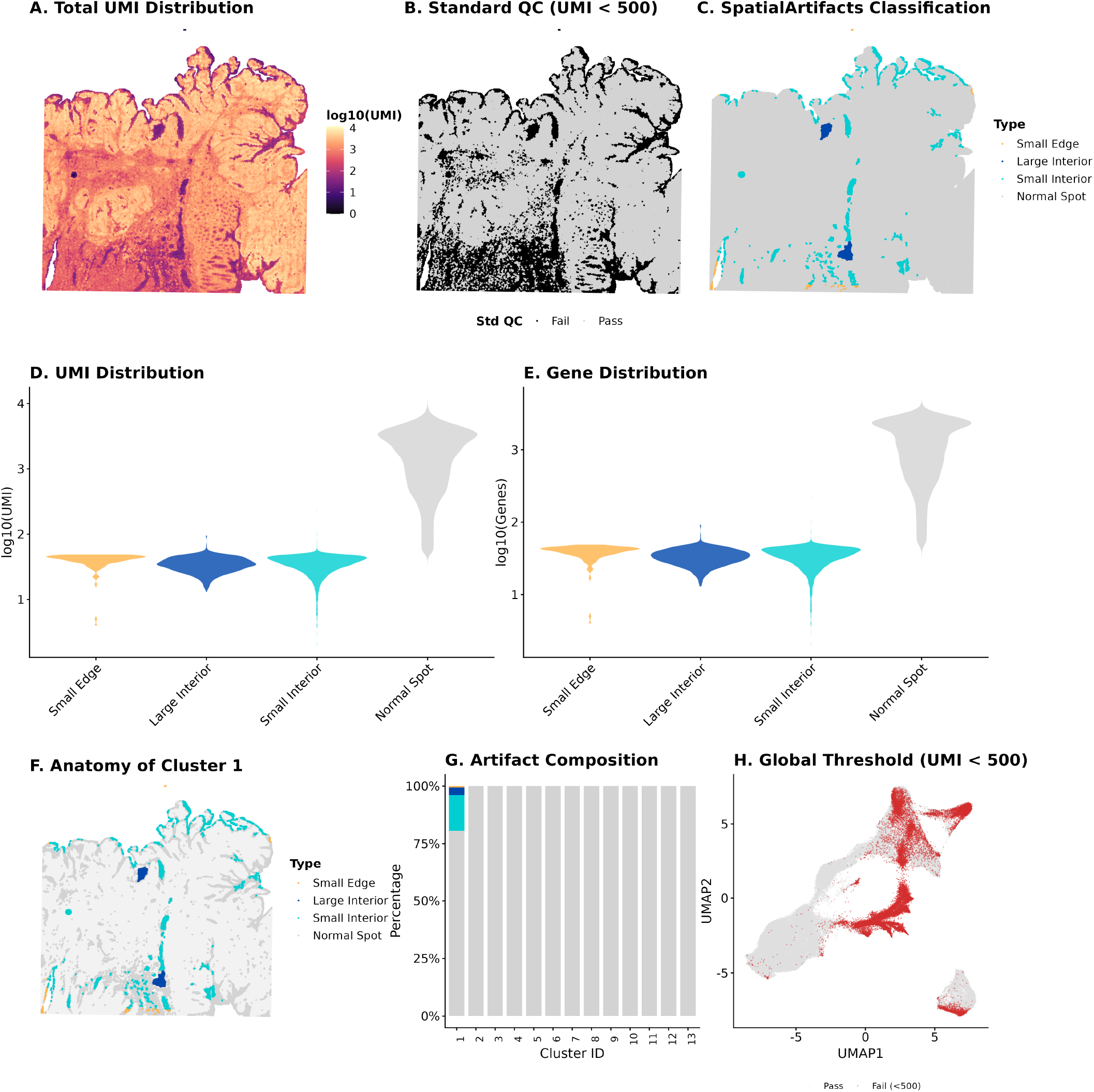
Computational scalability to high-resolution VisiumHD. Analysis of human colorectal cancer sample at 16-*µ*m resolution [4], demonstrating computational scalability moving from Visium (5,000 spots) to VisiumHD (137,051 bins). (**A**) Bin plot of total UMI counts (log_10_ scale) illustrates complex mucosal architecture. (**B**) Bin plot of low-quality bins (black) identified using standard QC thresholds (UMI *<* 500). (**C**) Bin plot with SpatialArtifacts classification including Small Edge, Large Interior, and Small Interior artifacts. (**D–E**) Violin plots confirming systematically reduced UMI counts (D) and gene detection (E) in detected artifacts, validating incomplete permeabilization at tissue boundaries. (**F**) Anatomical map of Cluster 1 showing spatial distribution of artifact bins at tissue edges and folds. (**G**) Artifact composition analysis reveals Cluster 1 contains 19.4% artifact bins (Small Edge, Large Interior, and Small Interior), while all other clusters consist predominantly of normal bins. (**H**) Global thresholding UMAP comparison (UMI *<* 500, red) shows massive indiscriminate removal including biologically coherent clusters, versus targeted removal by SpatialArtifacts.

Artifact compositional analysis of Cluster 1 (**Figure 4D-E**) revealed approximately 19.4% of the bins were artifacts (0.6% Small Edge, 3.3% Large Interior, 15.5% Small Interior), indicating technical rather than biological signal. Since only artifact bins within Cluster 1 were removed rather than the entire cluster, Cluster 1 is retained in the filtered data with its remaining 80.6% normal bins intact. SpatialArtifacts identified only 3,502 bins (2.6% of 137,050 total) as artifacts, preserving the underlying mucosal and muscularis tissue architecture. In contrast, global thresholding (UMI *<* 500) removed 31,688 bins (23.1%), of which 28,186 had a median UMI of 239 and were correctly preserved by SpatialArtifacts as biologically meaningful lowexpression regions (**Figure 4H, Figure S4E**). This nine-fold difference in removal rate demonstrates that SpatialArtifacts achieves targeted artifact removal without discarding biologically meaningful tissue regions.

We additionally validated the robustness of artifact detection using spot-level PCA. The variance structure remained stable before and after filtering (PC1: 16.98%→17.33%, PC2: 5.66%→5.31%), reflecting the biological specificity of SpatialArtifacts: since only 3,502 bins (2.6%) were identified as artifacts, the underlying mucosal and muscularis tissue architecture was preserved (**Figure S4C–D**). To confirm that the detected artifacts represent technical rather than biological signal loss, we examined the spatial expression of colorectal tissue marker genes EPCAM, ACTA2, and CDH1, which showed preserved expression in interior regions correctly retained by SpatialArtifacts (**Figure S4F–H**).

### 2.5 Method specificity validated through a negative control analysis

To evaluate the specificity of SpatialArtifacts, we selected a tissue section previously analyzed that had minimal tissue damage to evaluate the false-positive rate of classification. This tissue section profiled the gene expression from postmoretum human hippocampus [8]. We found that using a global threshold (total UMI *<*1000) (see **Figure S2G–I** for threshold comparison) resulted in a substantial number of spots being flagged, including regions known to have less UMIs (**Figure S5A-B**). In contrast, SpatialArtifacts detected very few edge artifacts, and all are also captured by the global threshold (**Figure S5C**). The remaining 618 spots flagged exclusively by the global threshold had a median UMI of 818, representing biologically meaningful low-expression regions correctly preserved by SpatialArtifacts (**Figure S6C**). Spatial clustering, UMAP, and QC distributions were nearly identical before and after filtering (**Figure S5F–K**), and compositional analysis revealed less than 1% artifact contamination across all clusters (**Figure S5H**).To further confirm that the detected artifacts represent technical rather than biological signal loss, we examined the spatial expression of hippocampal marker genes SNAP25 and GFAP, which showed preserved expression across the tissue interior, confirming the absence of significant technical damage in this sample (**Figure S6D–E**). Spot-level PCA confirmed that artifact classification explained minimal variance across all principal components, and the variance structure remained stable before and after filtering (**Figure S6A–B**). These results suggest that SpatialArtifacts is robust against false positives in tissues with high fidelity, a critical property for deployment across diverse datasets.

### 2.6 Benchmarking against existing methods demonstrates complementary QC approaches

We compared the low-quality spots identified from SpatialArtifacts to SpotSweeper [5] and BLADE [6] across three datasets: hippocampus (standard Visium, MAD = 2)[8], DLPFC (standard Visium, MAD = 3) [9] and Human colon (VisiumHD, MAD = 3) [11] (**Figure 5, Table S1**).

**Figure 5:**
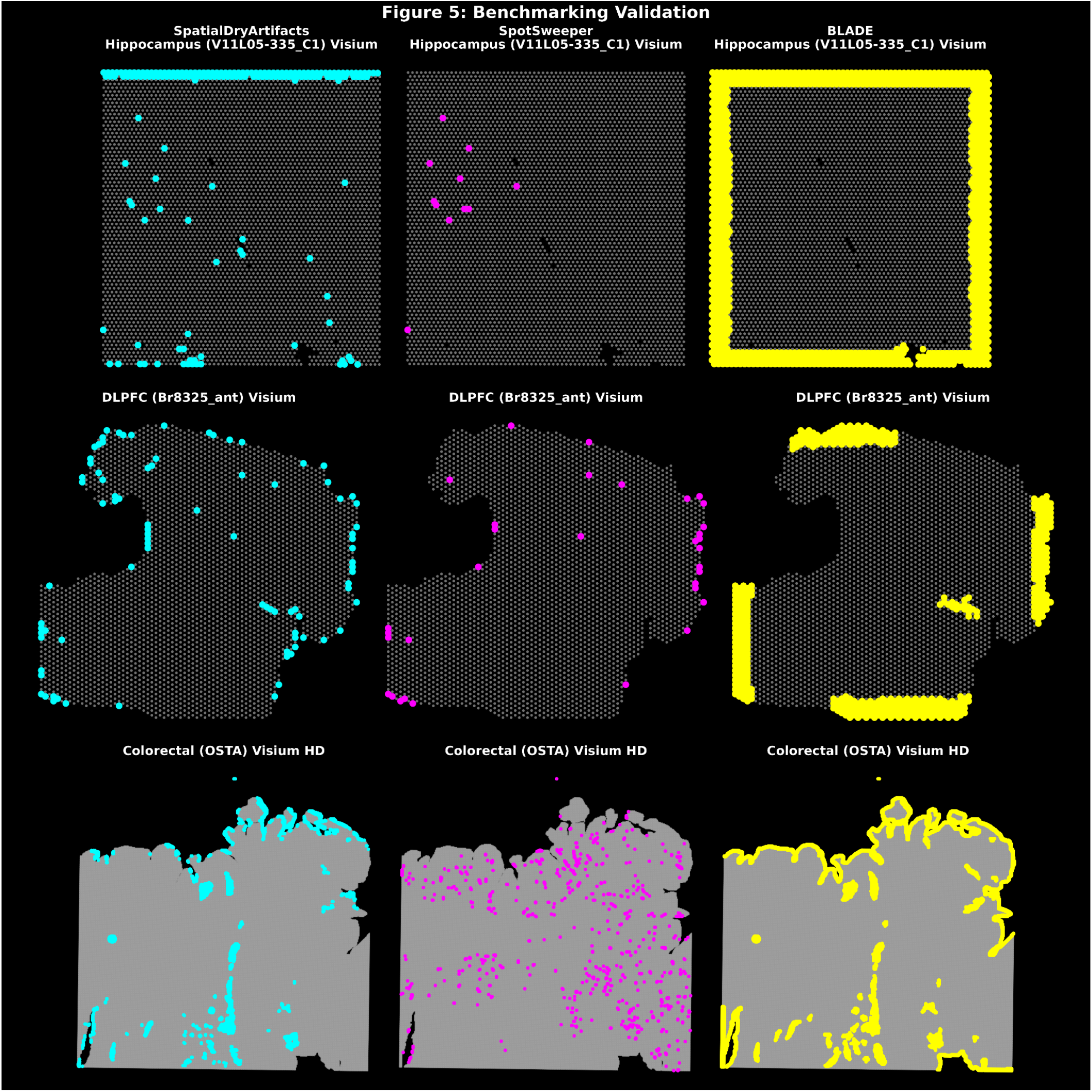
Benchmarking against existing methods demonstrates complementary detection strategies. Systematic comparison of SpatialArtifacts, SpotSweeper [5], and BLADE [6] across three representative datasets spanning platforms and tissue types. Rows represent datasets: Hippocampus (standard Visium, top)[8], DLPFC (standard Visium, middle) [9], and Human Colon (VisiumHD 16 *µ*m, bottom) [11]. Columns represent methods: SpatialArtifacts (left, cyan points), SpotSweeper (middle, magenta points), and BLADE (right, yellow regions). **Interpretation:** SpotSweeper excels at spot-level local outlier detection and biological heterogeneity normalization, while SpatialArtifacts targets spatially coherent regional artifacts through morphological reconstruction. Applied sequentially, these two tools offer layered quality control that balances sensitivity, specificity, and biological preservation. BLADE serves as a useful benchmark demonstrating the trade-off between simplicity and precision in edge artifact detection.

We found the three methods exhibited fundamentally different detection patterns. SpotSweeper consistently identified scattered individual spots based on *k*-nearest neighbor statistics, detecting only 11 spots (0.22%) in the hippocampus tissue. While it was effective at capturing low-quality spots associated certain synthetic DNA barcodes, the approach had limited sensitivity to larger spatially coherent artifacts. In contrast, BLADE removed all spots within a fixed distance from tissue boundaries, yielding substantially higher removal rates: 22.18% in the hippocampus, 13.06% in the DLPFC, and 5.60% in the VisiumHD colon. Its uniform ring removal lacks spot-level precision and risks discarding high-quality edge tissue. For example, spots located at the tissue boundary in the colon sample, but distant from the slide edge, were correctly classified by SpatialArtifacts as interior artifacts rather than edge artifacts, a distinction that BLADE’s fixed-distance approach fails to identify.

SpatialArtifacts detected 3-8× more artifacts than SpotSweeper, while removing 5-10× fewer spots than BLADE, detecting 167 spots (3.36%) in the hippocampus, 87 spots (2.47%) in the DLPFC (compared to SpotSweeper’s 33 spots (0.94%) and BLADE’s 461 spots (13.06%)), and 2,632 bins (1.92%) in the VisiumHD colon (compared to SpotSweeper’s 813 bins (0.59%) and BLADE’s 7,680 bins (5.60%)) (**Table S1**). Visual review confirmed that more than 95% of the removed spots exhibited low molecular capture and spatial coherence (cluster size *>* 20), supporting high precision. Together, these results suggest a complementary workflow: SpotSweeper for isolated low-quality spot removal followed by SpatialArtifacts for spatially coherently regional artifact detection, with BLADE serving as a useful benchmark for the trade-off between sensitivity and specificity in edge artifact detection.

## 3 Discussion

Spatial transcriptomics enables profiling of gene expression in its native tissue context [2], but downstream analyses depend critically on removing technical artifacts introduced during sample preparation [3, 4]. Here, we introduced SpatialArtifacts, which combines MAD-based outlier detection with mathematical morphology operations to precisely map the spatial footprint of tissue damage at a spot- or bin-level resolution in Visium and VisiumHD platforms.

SpatialArtifacts complements, rather than replaces, existing QC tools. SpotSweeper efectively detects stochastic dropouts due biases such as PCR amplification and preserves biologically valid low-expression domains using local QC metrics or a low variance in mitochondrial ratio for hangnails [5]. While this was an important step forward in spatially-aware QC, it can miss irregularly shaped edge artifacts. BLADE flags edge effects at the slide level, but it localize artifacts to individual spots [6], resulting in broad removal that risks discarding high-fidelity tissue. SpatialArtifacts filles this gap by providing spot-level artifact coordinates grouped into morphologically coherent clusters. We recommend applying these tools sequentially: SpotSweeper to remove isolated low-quality spots followed by SpatialArtifacts to resolve spatially coherent regional artifacts.

The primary technical contribution of SpatialArtifacts is the application of mathematical morphology operations, a methodology with roots in computer vision and medical imaging, to QC for spatial transcriptomics data. Edge artifacts share key properties with irregular lesions in MRI and histopathology images: their boundaries are shaped by physical properties such as surface tension and reagent flow rather than a predefined geometry which making morphological operations a natural fit [7]. The raster-based implementation via the terra package maintains the linear time complexity across both Visium and VisiumHD platforms [12], and the structural similarities between spatial transcriptomics data and geographic information systems suggest that the broader spatial data science toolkit could further advance QC methodology [13].

We also recognize several limitations with SpatialArtifacts. We have evaluated the performance on only two platforms, namely Visium and VisiumHD, and we have not evaluated this on non-grid platforms such as Slide-seqV2, Stereo-seq, or MERFISH. MAD-based thresholding is sensitive to the overall proportion of artifact-affected spots, and uses should manually adjust the `mad_threshold` parameter when artifact coverage exceeds approximately 20%. Third, there is no ground truth for edge artifacts, so detection accuracy is validated indirectly through improvements in downstream analyses such as unsupervised clustering. Curated benchmark datasets would help address this problem in the future. Finally, our current approches classifies artifacts based on location and size, but it does not describe a mechanism why an artifact occurred. This is left up to the analyst. Future work will include integrating H&E image features, selecting parameters in a more automated manner, and extensions to image-based platforms where negative control probes could serve as an alternative to quality signal.

## 4 Methods

### 4.1 Overview of SpatialArtifacts

We created an adaptive quality control framework for finding and identifying edge artifacts in spatial transcriptomics data based on mathematical morphology operations techniques. The framework has three key parts: (1) MAD-based outlier identification that works the same on all platforms, (2) platform-specific artifact detection algorithms, and (3) hierarchical classification of artifacts that have been found(**Figure 1**).

#### 4.1.1 Outlier identification via MAD thresholds

Using log_10_-transformed QC measures (library size or unique genes detected), we find low-quality regions for each tissue sample. Using the `isOutlier()` function from *scuttle* [14], we find outlier thresholds by applying a median absolute deviation (MAD) threshold of 3 (default) to each batch variable (slide, sample_id, or both).

Let *x*_*i*_ = log_10_(QC_metric_*i*_ + 1) for spot *i*. A spot is flagged as an outlier if:

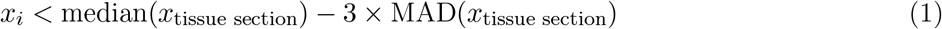

where the median and MAD are calculated over in-tissue spots within the same tissue section. It is important that to enable thorough artifact identification throughout the full capture area, the resulting thresholds are then applied to *all* spots, including background regions. This method guarantees that during the initial quality evaluation, edge artifacts that arise in regions with questionable tissue annotations are not overlooked. Subsequent platform-specific detection strategies use this first outlier flagging as input.

#### 4.1.2 Platform-specific detection strategies

In standard Visium data, edge artifacts are found by counting how many clusters of connected outliers fill a large part of the tissue borders that are described by the hexagonal `in_tissue` annotation. For VisiumHD data, the framework uses a physical border zone method instead of `in_tissue` flags, which are less reliable and have uneven tissue contours. Outliers within a defined buffer distance from the edge of the capture region are classified as edge artifacts (default: 80 *µ*m, automatically scaled to bin resolution). For interior problem area classification, clusters whose median coordinate falls within the buffer zone are reclassified as edge artifacts, while clusters centered in the tissue interior are retained as interior artifacts. Data-driven border detection can handle different tissue sizes while still being computationally efficient.

We use adaptive detection algorithms for each platform since standard Visium and VisiumHD platforms have very different grid architectures, tissue annotation reliability, and data scales (**Figure 1B-D**).

##### A. Standard Visium: morphological boundary analysis

We use mathematical morphology operations to connect geographically neighboring outliers and find edge artifacts based on biological tissue boundaries using standard Visium data (hexagonal grid, approximately 3,000–5,000 spots per sample with a 55 *µ*m spot diameter). Morphological operations on rasterized spatial data. Outlier spots are converted to binary raster format by constructing a `SpatRaster` grid via `terra::rast()` [15] based on the coordinate ranges of `array_row` and `array_col`, with outlier values assigned cell-by-cell. We apply a series of morphological transformations using focal operations to connect spatially adjacent outlier regions and remove isolated noise. The focal operations employ custom kernel functions implemented via `terra::focal()`:

1. **3**×**3 fill operation** (`my_fill()`): If all 8 of the surrounding locations are outliers, the center spot is recorded as an outlier for each spot in a 3×3 neighborhood. This operation fills in single-spot gaps within outlier regions.
2. **5**×**5 outline operation** (`my_outline()`): Examines a 5×5 window and marks the center point as an outlier if all 16 perimeter positions contain outliers. This captures larger-scale spatial patterns and connects nearby outlier clusters.
3. **Star-pattern fill** (`my_fill_star()`): Uses a star-shaped kernel considering only the four cardinal directions (north, south, east, and west). If all four cardinal neighbors are outliers, the center spot is marked as an outlier. This helps connect outlier regions along major axes while preserving diagonal boundaries.
4. **Small cluster removal**: After the fill operations, connected components of *non-outlier* spots are identified using `terra::patches()` with 8-directional connectivity and `zeroAsNA=TRUE` to treat background pixels (value = 0) identically to NA values. Isolated non-outlier regions smaller than `min_cluster_size` (default: 40 spots) are converted to outliers, effectively removing noise from the cleaned outlier map.

These operations are implemented using the `terra::focal()` function with custom convolution kernels and the `na.policy=“all”` parameter to handle edge effects. For hexagonal grid arrangements, an optional coordinate adjustment (`shifted = TRUE`) can be applied to account for the offset pattern of odd-numbered columns.

##### Edge detection via dual-strategy boundary analysis

After morphological cleaning, connected components of outlier spots are identified using `terra::patches()` with 8-directional connectivity and `zeroAsNA=TRUE`, ensuring proper separation of outlier clusters from background. For each identified cluster, we determine whether it represents an edge artifact using a dual-strategy approach that combines coverage-based and touch-based detection. Clusters identified by either method are classified as edge artifacts, ensuring both large continuous dryspots and smaller scattered edge artifacts are captured.

##### Coverage-based detection

For each of the four border directions (north, south, east, and west), we calculate:

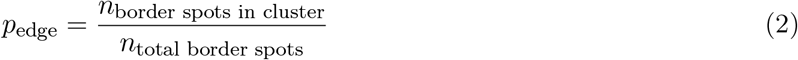

where *n*_border spots in cluster_ is the number of spots along that border belonging to the cluster and *n*_total border spots_ is the total number of in-tissue spots along that border. A cluster is classified as an edge artifact if *p*_edge_ ≥ `edge_threshold` (default: 0.75) for any single border direction.

##### Touch-based detection

To capture scattered edge artifacts that may not exceed the coverage criterion, we further identify every cluster that has any contact with the capture area boundaries of the raster grid (regardless of coverage percentage) as an edge artifact. Clusters identified by either method are classified as edge artifacts, ensuring both large continuous edge dryspots and smaller scattered artifacts along tissue margins are captured.

##### Interior problem area detection

All connected outlier clusters, regardless of edge status, are identified and characterized. Each cluster receives a unique identifier (formatted as `{sample_id}_{cluster_number}`) and size annotation (total number of spots), enabling comprehensive quality tracking across samples and flexible downstream filtering decisions.

##### B. VisiumHD: Physical boundary zone approach

VisiumHD data poses distinct challenges that require an alternative detection strategy: (1) a square grid architecture with enhanced spatial resolution (8–16 *µ*m bins compared to 55 *µ*m spots), resulting in 10–100 times more observations per sample; (2) irregular tissue contours that hinder biological boundary detection; and (3) less reliable `in_tissue` annotations due to automated segmentation applied at scale to dense bin data. We use a physical border zone technique that takes advantage of the fixed sizes of VisiumHD capture regions and resolution.

##### Resolution-aware parameter specification

To ensure consistent detection across different VisiumHD resolutions (8 *µ*m vs. 16 *µ*m), we specify detection parameters in physical units rather than bin counts. The default values for the buffer zone width and minimum cluster size are 80 *µ*m and 1,280 *µ*m^2^, respectively. These values are automatically converted to bin-space equivalents based on the specified resolution:

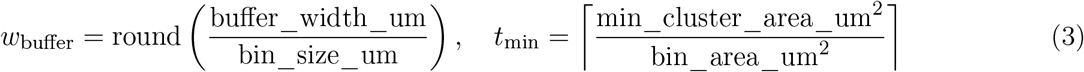

where bin_size_um ∈ {8, 16} *µ*m and bin_area_um^2^ = bin_size_um^2^ ∈ {64, 256} *µ*m^2^. For example, with default parameters at 16 *µ*m resolution: *w*_buffer_ = round(80*/*16) = 5 bins and *t*_min_ = ⌈1280*/*256⌉ = 5 bins. At 8 *µ*m resolution, these become 10 bins and 20 bins respectively, maintaining equivalent physical coverage.

##### Edge detection via data-driven physical boundaries

Instead of using predetermined capture area sizes, we extract actual data boundaries from the observed coordinate ranges to accommodate tissue-specific variations. For VisiumHD data, edge artifacts are identified as outliers whose bin coordinates (*x*_*i*_, *y*_*i*_) satisfy:

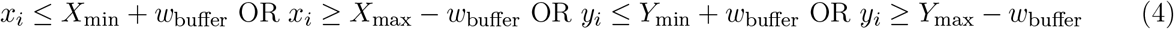

where *X*_min_, *X*_max_, *Y*_min_, and *Y*_max_ are extracted from the actual data range via `range(colData(spe)`$`array_col)` and `range(colData(spe)`$`array_row)`, and *w*_buffer_ is the resolutionadjusted buffer width in bins. A bin is classified as an edge artifact if it satisfies the above spatial condition *and* was identified as a MAD-based outlier in Stage 1. This data-driven approach accommodates variable tissue sizes and capture area utilization while maintaining computational efficiency (*O*(*n*) complexity, where *n* is the number of bins).

The physical boundary zone approach is motivated by the mechanism of edge dryspot formation: incomplete reagent coverage during permeabilization affects regions near the physical edges of the capture area, independent of the shape of the tissue section. By focusing on physical rather than biological boundaries, we avoid confounding from irregular tissue contours and segmentation artifacts while maintaining high sensitivity for true edge artifacts.

##### Interior problem area detection via terra-optimized morphological clustering

For outliers not classified as edge artifacts, we apply morphological processing comparable to standard Visium, but optimized for large-scale VisiumHD datasets using the *terra* package [15]. The morphological transformation pipeline (`focal_transformations()`) applies the same sequence of operations adapted for square grid architecture: 3×3 fill, 5×5 outline, and star-pattern operations.

Connected outlier regions are identified using `terra::patches()` with 8-directional connectivity and `zeroAsNA=TRUE` to ensure proper cluster separation. To leverage terra’s performance optimizations for large rasters, we use memory-efficient bin-based coordinates (`array_col`, `array_row`) rather than pixel-based coordinates, reducing memory requirements by 10–100× for typical VisiumHD datasets.

Detected clusters are further filtered by their median coordinate: clusters whose center falls within the buffer zone are reclassified as edge artifacts, while those centered in the tissue interior are retained as interior problem areas. The resolution-aware minimum cluster size threshold (*t*_min_, automatically scaled from physical units as described above) prevents over-aggressive removal of small but meaningful outlier clusters while adapting to the higher spatial resolution of VisiumHD data.

#### 4.1.3 Hierarchical classification scheme

Detected artifacts are organized into a four-tier hierarchy depending on their spatial location (edge vs. interior) and cluster size. The unified `classifyEdgeArtifacts()` function does this for both standard Visium and VisiumHD data (**Figure 1F–H**). This hierarchical architecture enables flexible downstream filtering strategies adapted to different biological contexts and quality control stringencies.

##### Classification logic

Let *C* be a detected cluster with size |*C*| spots (or bins for VisiumHD). Classification proceeds hierarchically with edge location taking priority:

1. **Edge artifact determination:** If cluster *C* was flagged as an edge artifact (via dual-strategy boundary analysis for standard Visium, or physical buffer zones for Visium HD), proceed to size classification as an edge artifact. Otherwise, classify as an interior artifact and proceed to size classification.
2. **Size-based sub-classification:** For each artifact type (edge or interior), further classify by cluster size:

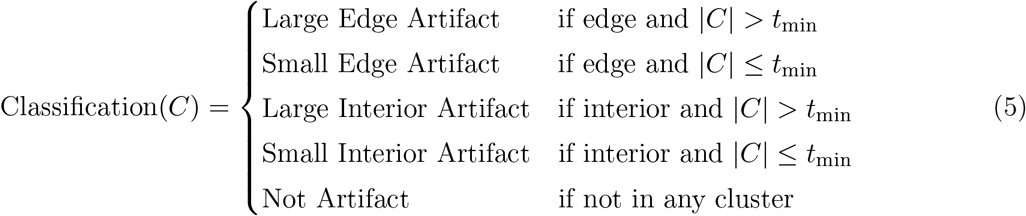

where *t*_min_ is the minimum cluster size threshold for classification (default: 20 spots for standard Visium; 100–200 bins for VisiumHD at 16 *µ*m resolution and 400–800 bins at 8 *µ*m resolution, scaled to maintain equivalent physical artifact area).

##### Slide-specific exclusions

The optional `exclude_slides` argument allows masking edge artifacts from specific slides with known technical issues unrelated to edge dryspots (e.g., unusual tissue placement or capture area manufacturing defects). This creates a `true_edges` indicator that distinguishes edge artifacts from slide-specific artifacts, enabling more accurate downstream quality metrics.

##### Classification outputs

The complete pipeline generates five complementary `colData` columns in the `SpatialExperiment` object:

- `edge_artifact_classification`: Five-level hierarchical classification (large/small × edge/interior, plus `not_artifact`)
- `edge_artifact_edge`: Binary edge artifact indicator
- `edge_artifact_true_edges`: Edge indicator with slide-specific exclusions applied
- `edge_artifact_problem_id`: Unique cluster identifier for all detected problem areas
- `edge_artifact_problem_size`: Cluster size (number of spots/bins)

This hierarchical scheme enables flexible downstream filtering strategies. For example, analysts can remove all edge artifacts while retaining large interior artifacts for careful manual review, or apply different stringency thresholds based on tissue type and experimental goals. The cluster size information allows assessment of whether problem areas represent systematic artifacts or isolated technical failures.

### 4.2 Summary of datasets

#### 4.2.1 Human hippocampus on the 10x Genomics Visium platform

We analyzed 36 Visium spatial transcriptomics samples of postmortem human hippocampus tissue from Thompson et al. [8]. Two representative samples were selected for detailed visualization: sample `V11L05-335_C1` for primary artifact detection analysis, and sample `V11U08-081_B1` as a negative control to assess method specificity. For each sample, standard quality control metrics were calculated using `addPerCellQCMetrics()` from *scuttle* [14], including:

- Library size (total UMI counts per spot)
- Number of unique genes detected
- Mitochondrial read percentage (not used for edge detection)

#### 4.2.2 Human DLPFC on the 10x Genomics Visium platform

We analyzed sample `Br8325_ant` from the human dorsolateral prefrontal cortex (DLPFC) dataset from Huuki-Myers et al. [9], which includes manual annotations of six cortical layers and white matter. These annotations served as ground truth for evaluating artifact removal and cortical layer delineation.

#### 4.2.3 Human Colorectal Cancer on the 10x Genomics VisiumHD platform

We analyzed a human colorectal cancer sample at 16 *µ*m resolution [11], generated on the 10x Genomics VisiumHD platform [4]. Standard quality control metrics were calculated using `addPerCellQCMetrics()` from *scuttle* [14].

### 4.3 Details related to the SpatialArtifacts software package

Full implementation details, default parameters, and usage tutorials are available in the R/Bioconductor package vignette at https://bioconductor.org/packages/SpatialArtifacts and in the Python package tutorials on GitHub at https://github.com/CambridgeCat13/SpatialArtifacts-py or access through PyPI https://pypi.org/project/spatial-artifacts/.

### 4.4 Summary of exisiting QC methods

To demonstrate the performance and added value of SpatialArtifacts, we compare our approach to other state-of-the-art tools for quality control in spatial transcriptomics, in particular SpotSweeper [5] and BLADE [6]. Although all of these methods are intended to reveal and mitigate the effects of technical artifacts, they differ fundamentally in philosophical approach, the specific types of artifacts they target, and the type of output they produce.

#### 4.4.1 Comparison with SpotSweeper

SpotSweeper [5] brings a “spatially aware” QC scheme, aimed originally at countering biases emanating from biological heterogeneity in a tissue section. Fundamental to the success of this work is a locally normalized representation.

- Spot-level QC: Low-quality spots are detected individually by computing a robust z-score for each spot in comparison to its neighboring spots (k-nearest neighbors). This is in sharp contrast to our approach, which begins by identifying outliers across the entire batch (e.g., slide or sample) to establish a global quality baseline before employing spatial information for grouping. SpotSweeper’s approach is particularly effective at avoiding the inadvertent removal of spots in biologically low-expressing areas, such as white matter regions in the brain [5], which would be flagged by a naively chosen global threshold.
- Region-level artifacts: SpotSweeper defines and targets two specific types of regional artifacts: “dryspots,” found by clustering over QC metrics especially library size, and “hangnails” (tissue damage), which are identified by finding regions with abnormally low variance in mitochondrial expression.

`SpatialArtifacts` instead uses a mathematical morphology operation approach. Rather than hand-designing artifact signatures, it takes a bottom-up approach: flagging all low-quality spots and then using mathematical morphology operations to determine if they are spatially clustering. The robustness of our approach stems from a geometrically defined algorithm for identifying edge artifacts by calculating their proportional coverage of the tissue boundary, a role not explicitly performed within SpotSweeper.

#### 4.4.2 Comparison with BLADE

BLADE [6] (Border, Location, and edge Artifact Detection) is a statistical approach for detecting three pre-specified types of artifacts: tissue edge effects, capture area border effects, and location batch malfunctions.

- **Methodological approach:** BLADE operates by defining spatial zones based on taxicab distance from the tissue edge or capture area border, and then applying a two-sample unpaired t-test to compare the distribution of gene read counts between those zones and interior spots.
- **Output:** The output of this analysis is a slide-level P-value indicating whether the entire slide exhibits an edge or border effect. This provides a statistical measure at the sample level, but does not identify which specific spots constitute the artifact region.

In contrast, SpatialArtifacts does not perform a statistical hypothesis test for the presence of an artifact. Rather, it operates at the spot level, assigning each spot to a spatially coherent artifact cluster. Consequently, whereas BLADE addresses the question “Is there an edge artifact on this slide?”, our package addresses the more refined question “Which exact spots constitute the edge artifact on this slide?” Our mathematical morphology-based clustering algorithm was specifically developed to delineate these artifacts at spot-level resolution for targeted removal or masking.

## 4.5 Code and software availability

Code to reproduce all preprocessing, analyses, and figures in this manuscript is available on GitHub at https://github.com/CambridgeCat13/SpatialArtifacts–paper. SpatialArtifacts is available as an open-source R/Bioconductor package at https://bioconductor.org/packages/SpatialArtifacts and on GitHub at https://github.com/CambridgeCat13/SpatialArtifacts. A Python implementation is available on PyPI at https://pypi.org/project/spatial-artifacts/ and on GitHub at https://github.com/CambridgeCat13/SpatialArtifacts-py.

## 4.6 Acknowledgments, Funding, Authorship Contributions

## Acknowledgments

We acknowledge the Joint High Performance Computing Exchange (JHPCE) at the Johns Hopkins Bloomberg School of Public Health for computational support, and members of the Hicks lab for their valuable feedback.

## Author Contributions Statement

**JHH:** Software, Data curation, Methodology, Formal analysis, Investigation, Visualization, Writing – original draft, Writing – review & editing; **JRT:** Conceptualization, Methodology, Software, Formal analysis, Supervision, Writing – review & editing; **MT:** Supervision, Writing – review & editing; **SCH:** Supervision, Funding acquisition, Project administration, Software, Writing – original draft, Writing – review & editing.

## Supplementary Materials

**Table S1:** Quantitative benchmarking of artifact detection methods across platforms and tissue types.

| Dataset | Platform | Method | Total Spots | Artifacts | % Removed |
| --- | --- | --- | --- | --- | --- |
| Hippocampus | Standard | BLADE | 4,965 | 1,101 | 22.18% |
|  |  | SpotSweeper | 4,965 | 11 | 0.22% |
|  |  | **SpatialArtifacts** | 4,965 | **167** | **3.36%** |
| DLPFC | Standard | BLADE | 3,529 | 461 | 13.06% |
|  |  | SpotSweeper | 3,529 | 33 | 0.94% |
|  |  | **SpatialArtifacts** | 3,529 | **87** | **2.47%** |
| VisiumHD (16μm) | HD | BLADE | 137,051 | 7,680 | 5.60% |
|  |  | SpotSweeper | 137,051 | 813 | 0.59% |
|  |  | **SpatialArtifacts** | 137,051 | **2,632** | **1.92%** |

**Figure S1:**
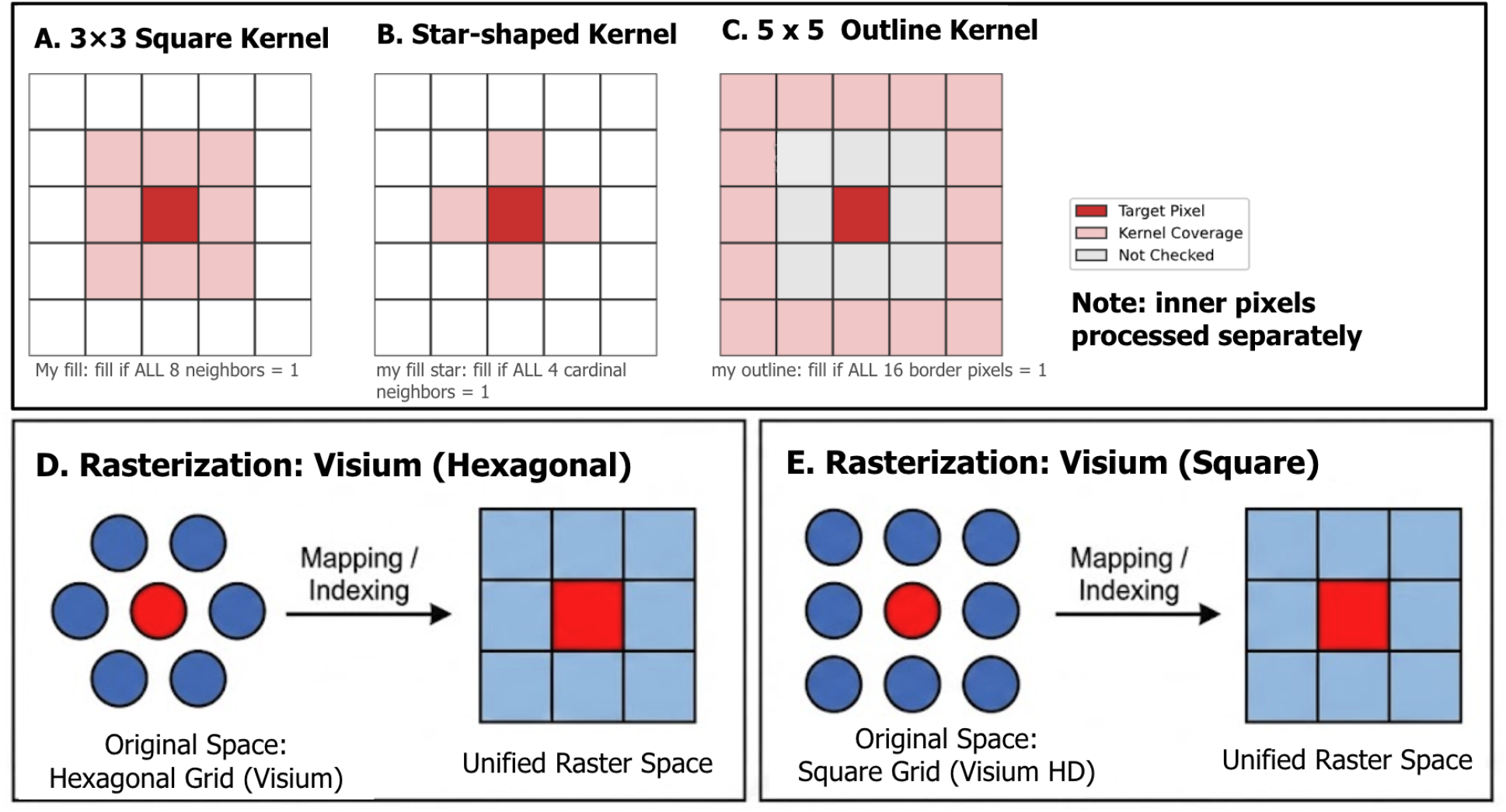
Overview of computational approaches for morphological pattern recognition. (**A**-**C**) Schematic of the focal filling kernels used for artifact reconstruction, including the 3 × 3 square kernel, star-shaped cardinal neighbor kernel, and 5 × 5 outline kernel. (**D**-**E**) Illustration of the rasterization process used to map hexagonal (standard Visium) and square (VisiumHD) spot coordinates into a unified raster space for morphological processing.

**Figure S2:**
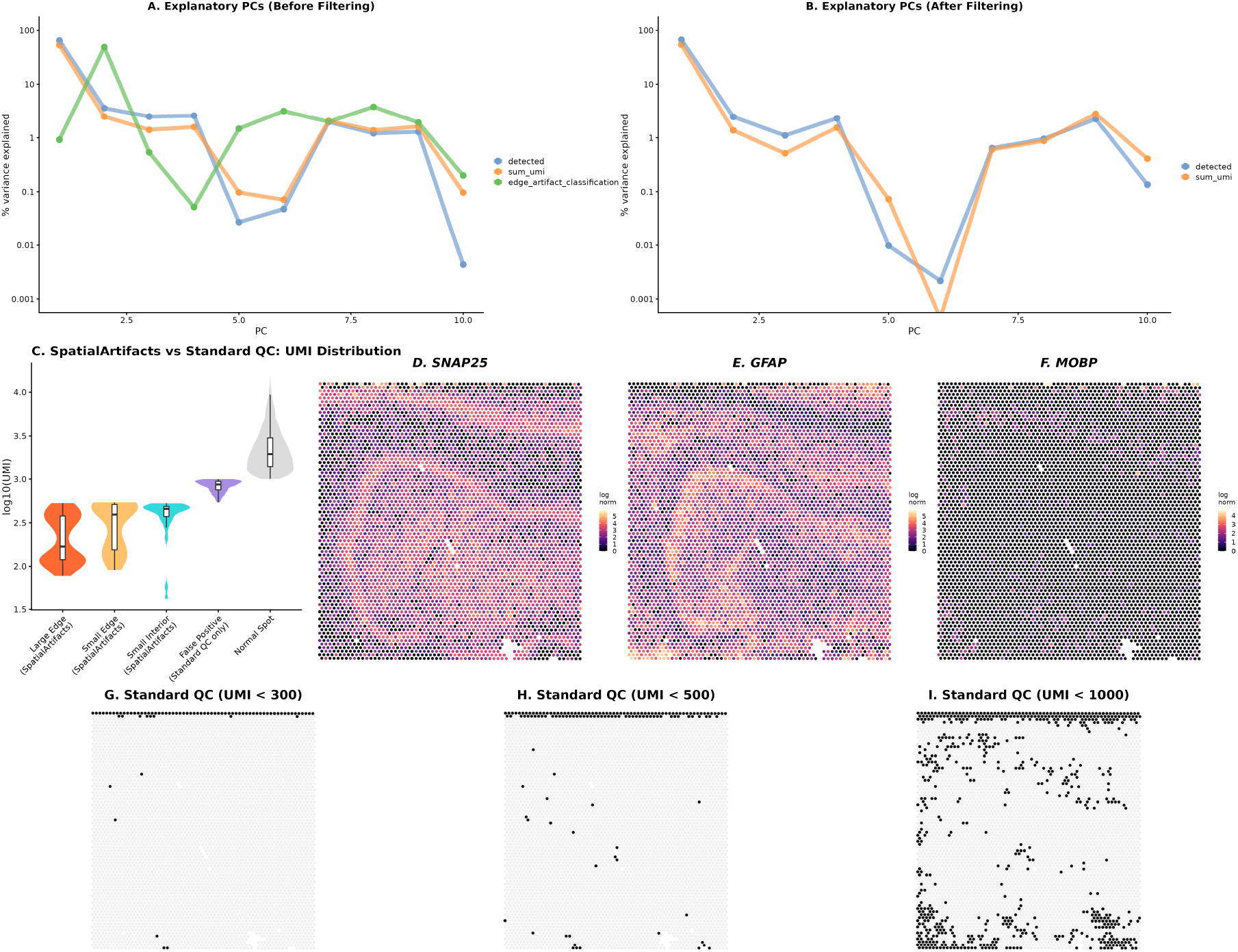
Spot-level PCA, QC validation, and threshold comparison for human hippocampus sample V11L05-335_C1. (**A–B**) Assessment of metadata variable contributions to the top 10 principal components using `plotExplanatoryPCs`, before (A) and after (B) artifact removal. Prior to filtering, artifact classification explained a substantial proportion of variance in PC1 and PC2. After removing artifact spots, the contribution of QC metrics to mid-range PCs (PC4–6) decreased by approximately 100-fold, suggesting that these components previously captured artifact-driven technical variation.(**C**) UMI distribution comparison between SpatialArtifacts artifact categories and spots flagged exclusively by standard global thresholding (UMI *<* 1000). The 492 false positive spots flagged only by standard QC show a median UMI of 840, consistent with biologically meaningful low-expression regions rather than technical artifacts.(**D–F**) Spatial expression of hippocampal marker genes SNAP25 (D), GFAP (E), and MOBP (F), confirming preserved biological signal in interior regions with naturally lower UMI counts.(**G–I**) Comparison of three fixed QC thresholds (UMI *<* 300, UMI *<* 500, and UMI *<* 1000).

**Figure S3:**
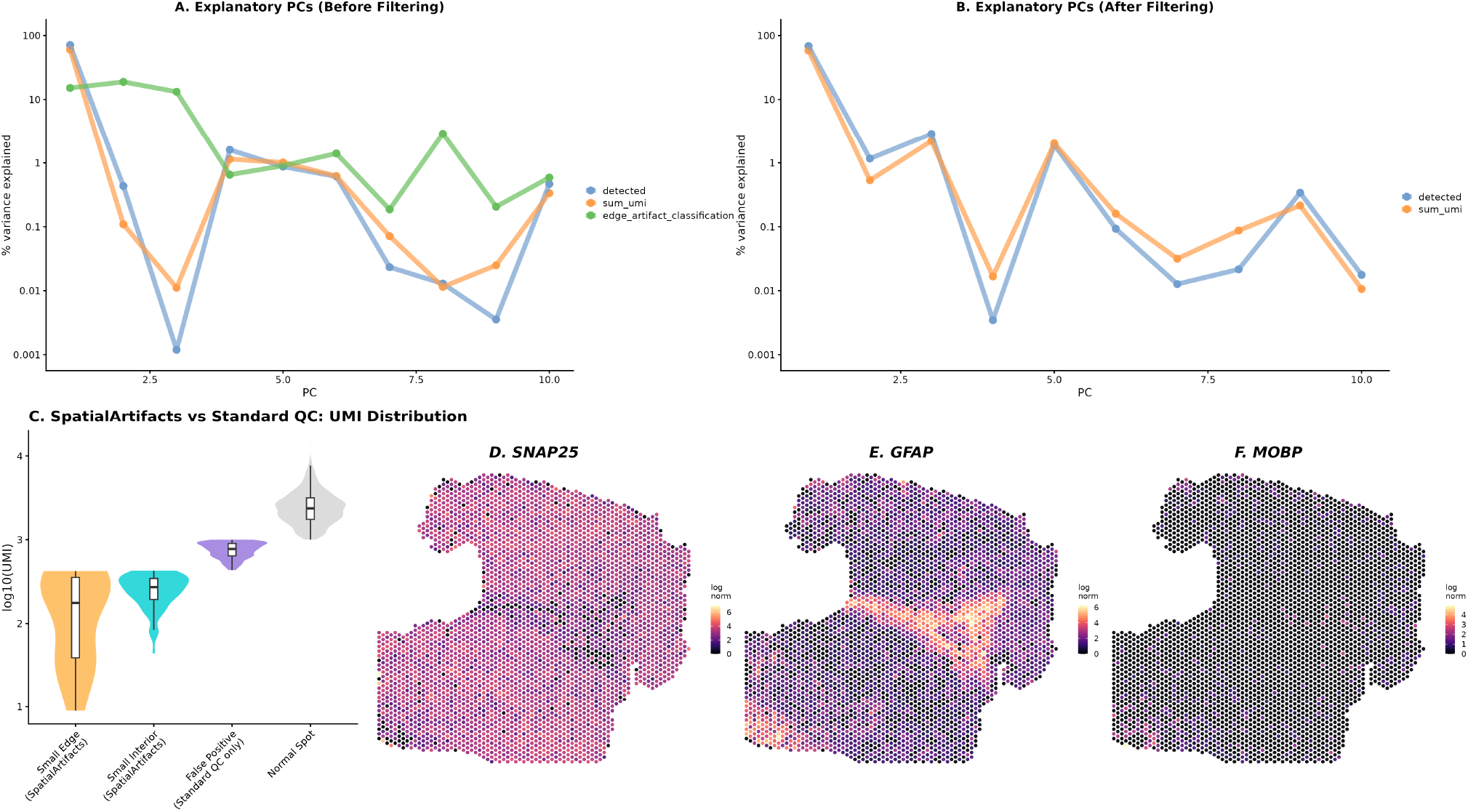
Spot-level PCA and QC validation for DLPFC sample Br8325_ant. (**A–B**) Assessment of metadata variable contributions to the top 10 principal components using `plotExplanatoryPCs`, before (A) and after (B) artifact removal. Prior to filtering, artifact classification explained a substantial proportion of variance in PC1 and PC2. After removal, the contribution of QC metrics was reduced, suggesting that these components previously captured artifact-driven technical variation.(**C**) UMI distribution comparison between SpatialArtifacts artifact categories and spots flagged exclusively by standard global thresholding (UMI *<* 1000). The 389 false positive spots flagged only by standard QC show a median UMI of 773, consistent with biologically meaningful low-expression regions such as white matter.(**D–F**) Spatial expression of cortical marker genes SNAP25 (D), GFAP (E), and MOBP (F). Artifact spots retained partial SNAP25 expression (median = 2.84) but showed complete loss of GFAP expression (median = 0), consistent with selective technical degradation rather than complete signal loss.

**Figure S4:**
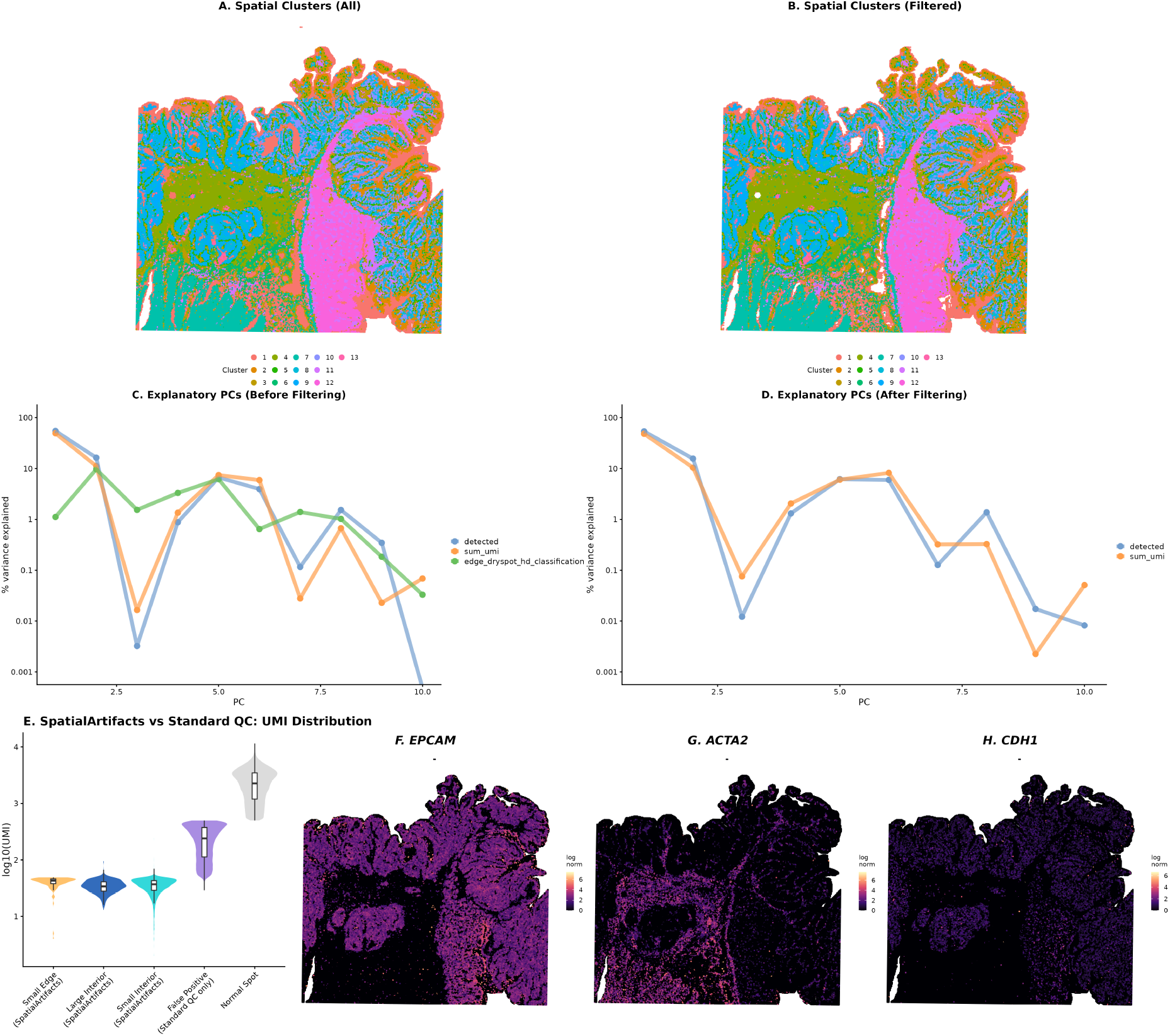
Comprehensive validation on high-resolution VisiumHD colorectal cancer data. (**A– B**) Spatial cluster maps before (A) and after (B) artifact removal, showing improved domain detection with all clusters retained.(**C–D**) Assessment of metadata variable contributions to the top 10 principal components using `plotExplanatoryPCs`, before (C) and after (D) artifact removal. Prior to filtering, artifact classification explained a substantial proportion of variance; after removal, QC metric contributions were reduced, confirming biological specificity of artifact detection. (**E**) UMI distribution comparison between SpatialArtifacts artifact categories and bins flagged exclusively by standard global thresholding (UMI *<* 500). The 28,186 false positive bins flagged only by standard QC show a median UMI of 239, consistent with biologically meaningful low-expression regions rather than technical artifacts.(**F–H**) Spatial expression of colorectal tissue marker genes EPCAM (F), ACTA2 (G), and CDH1 (H), confirming preserved biological signal in regions with naturally lower UMI counts that are correctly retained by SpatialArtifacts.

**Figure S5:**
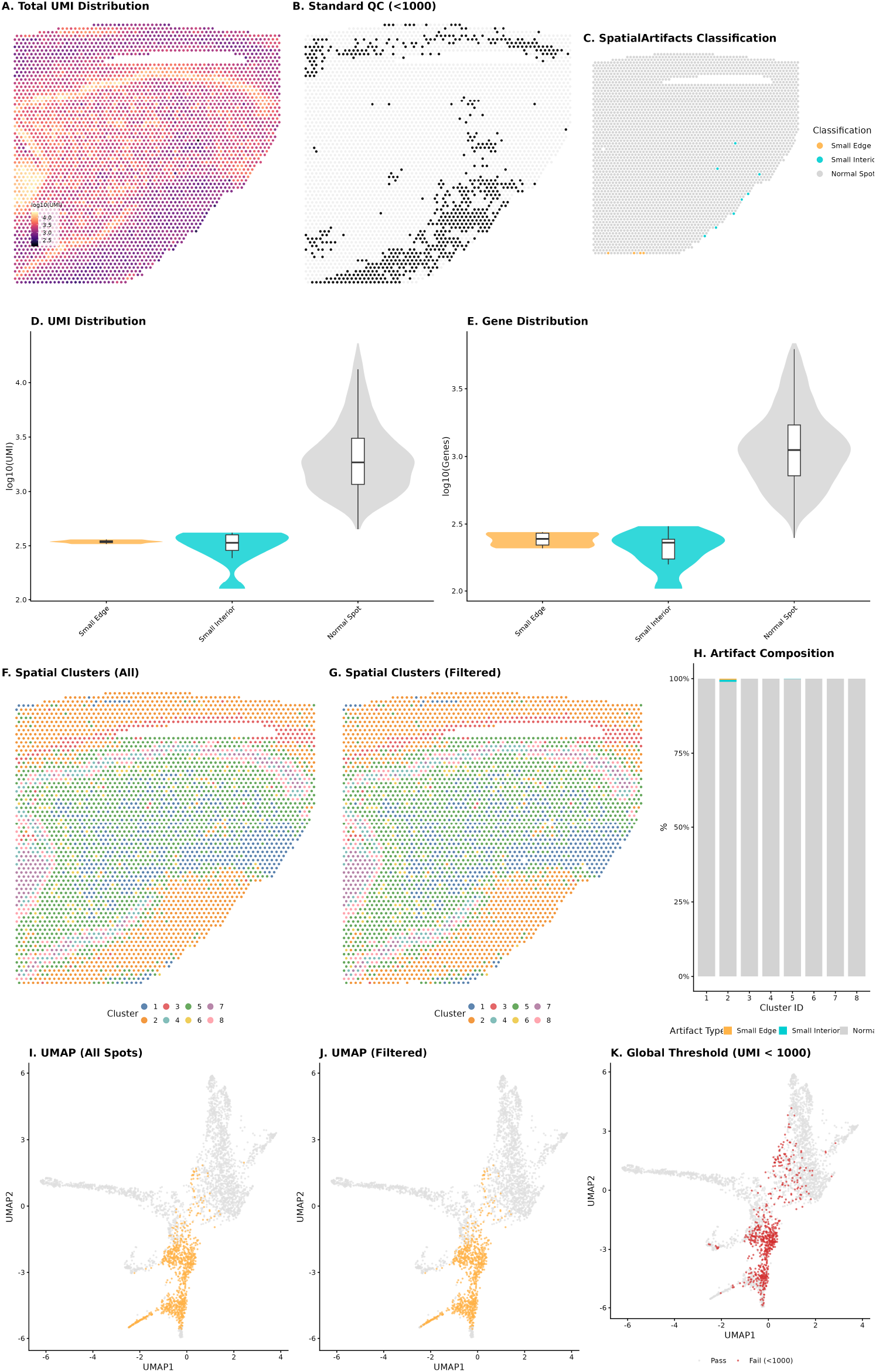
Negative control analysis using a high-quality human hippocampus sample. Evaluation of Sample V11U08-081_B1, which exhibits minimal technical damage. (**A**) Heat map of total UMI (log_10_ scale). (**B**) Spots flagged as low-quality (black) using standard global QC thresholds (UMI *<* 1000). (**C**) SpatialArtifacts classification confirming detection of only 12 artifact spots (4 Small Edge and 8 Small Interior). (**D–E**) Violin plots for the distribution of UMI counts (D) and number of detected genes (E). (**F–G**) Spatial clustering before (F) and after (G) artifact removal, demonstrating maintenance of biological structure with all 8 clusters retained. (**H**) Artifact composition analysis revealing less than 1% artifact contamination across all clusters. (**I–J**) UMAPs before (I) and after (J) filtering, confirming no spurious cluster removal. (**K**) Global thresholding comparison (UMI *<* 1000, red points) removes substantially more spots than SpatialArtifacts.

**Figure S6:**
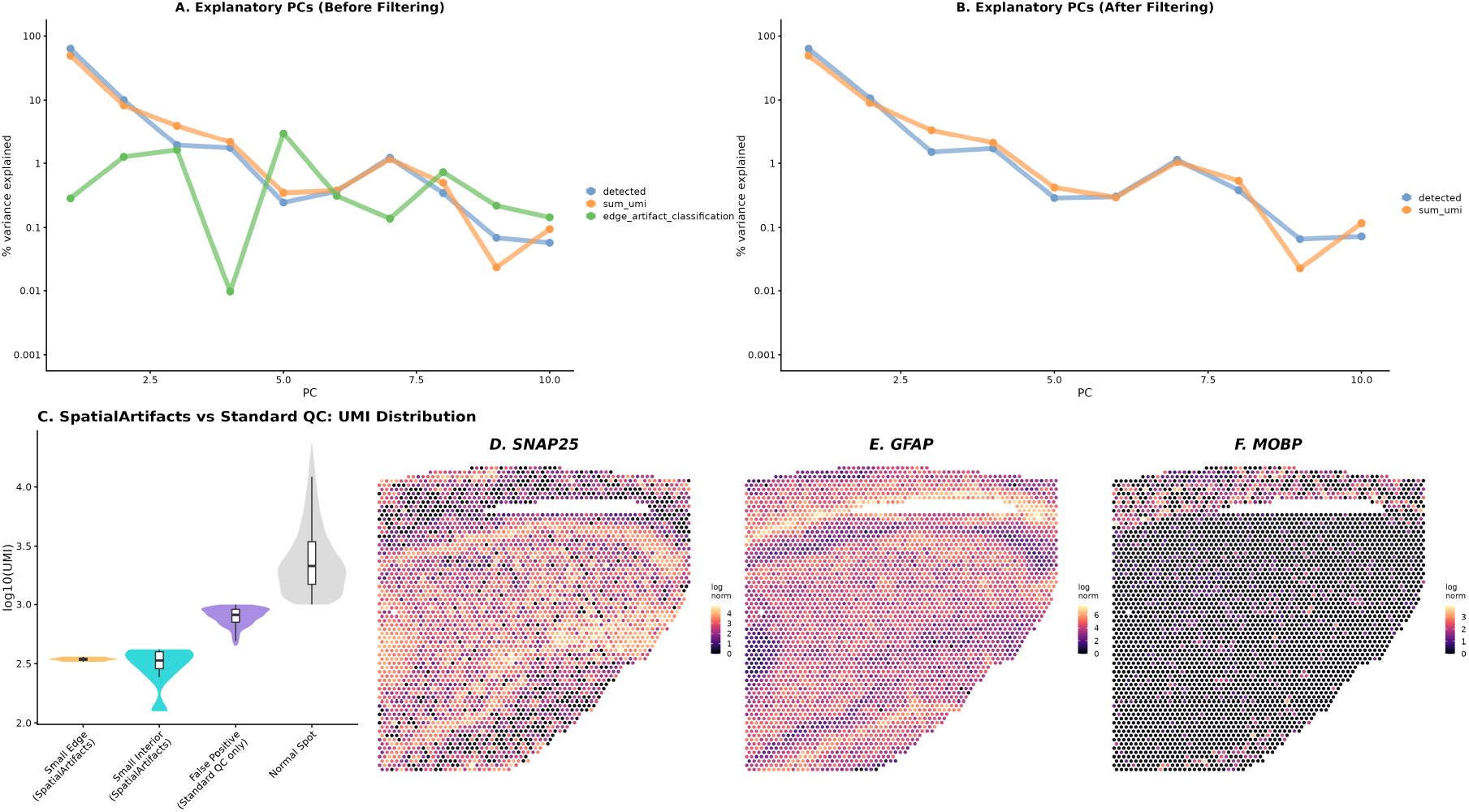
Spot-level PCA and QC validation for negative control Sample V11U08-081_B1. (**A– B**) Assessment of metadata variable contributions to the top 10 principal components using `plotExplanatoryPCs`, before (A) and after (B) artifact removal. The variance structure remains stable, with artifact classification explaining minimal variance, confirming that SpatialArtifacts does not remove biologically meaningful variation in high-quality tissue. (**C**) UMI distribution comparison between SpatialArtifacts artifact categories and spots flagged exclusively by standard global thresholding (UMI *<* 1000). The 618 false positive spots flagged only by standard QC show a median UMI of 818, consistent with biologically meaningful low-expression regions rather than technical artifacts. (**D–F**) Spatial expression of hippocampal marker genes SNAP25 (D), GFAP (E), and MOBP (F), showing preserved expression across the tissue interior, further confirming the absence of significant technical damage in this sample.

## References

[1] P. L. Ståhl, F. Salmén, S. Vickovic, A. Lundmark, J. F. Navarro, J. Magnusson, S. Giacomello, M. Asp, J. O. Westholm, M. Huss, A. Mollbrink, S. Linnarsson, S. Codeluppi, Borg, F. Pontén, P. I. Costea, P. Sahlén, J. Mulder, O. Bergmann, J. Lundeberg, and J. Frisén. Visualization and analysis of gene expression in tissue sections by spatial transcriptomics. Science (New York, N.Y.), 353(6294):78–82, July 2016. ISSN 1095-9203. doi:10.1126/science.aaf2403.

[2] V. Marx. Method of the Year: spatially resolved transcriptomics. Nature Methods, 18(1):9–14, Jan. 2021. ISSN 1548-7091, 1548-7105. doi:10.1038/s41592-020-01033-y. URL https://www.nature.com/articles/s41592-020-01033-y.

[3] 10x Genomics. Visium Spatial Gene Expression Reagent Kits User Guide. Technical Report CG000239 Rev H, 10x Genomics, 2024. URL https://cdn.10xgenomics.com/image/upload/v1723742999/support-documents/CG000239_Visium_Spatial_Gene_Expression_User_Guide_RevH.pdf.

[4] 10x Genomics. Visium HD FFPE Tissue Preparation Handbook. Technical Report CG000684 Rev A, 10x Genomics, 2024. URL https://cdn.10xgenomics.com/image/upload/v1711129979/CG000684_VisiumHDFFPETissuePrepHandbook_RevA.pdf.

[5] M. Totty, S. C. Hicks, and B. Guo. SpotSweeper: spatially aware quality control for spatial transcriptomics. Nature Methods, 22(7):1520–1530, July 2025. ISSN 1548-7091, 1548-7105. doi:10.1038/s41592-025-02713-3. URL https://www.nature.com/articles/s41592-025-02713-3.

[6] E. Kummerfeld, L. Williams, Y. Wang, S. T. Peters, E. Schmidt, M. DuFresne-To, D. Bernlohr, P. Robbins, S. Ikramuddin, O. Adeyi, L. Laux, G. Barthel, S. Johnson, J. Wang, L. Niedernhofer, A. Nelson, and C. Aliferis. Artifacts in spatial transcriptomics data: their detection, importance, prevalence, and prevention. Briefings in Bioinformatics, 26(4):bbaf306, July 2025. ISSN 1477-4054. doi:10.1093/bib/bbaf306.

[7] R. M. Haralick, S. R. Sternberg, and X. Zhuang. Image Analysis Using Mathematical Morphology. IEEE Transactions on Pattern Analysis and Machine Intelligence, PAMI-9(4):532–550, July 1987. ISSN 0162-8828. doi:10.1109/TPAMI.1987.4767941. URL https://ieeexplore.ieee.org/document/4767941.

[8] J. R. Thompson, E. D. Nelson, M. Tippani, A. D. Ramnauth, H. R. Divecha, R. A. Miller, N. J. Eagles, E. A. Pattie, S. H. Kwon, S. V. Bach, U. M. Kaipa, J. Yao, C. Hou, J. E. Kleinman, L. Collado-Torres, S. Han, K. R. Maynard, T. M. Hyde, K. Martinowich, S. C. Page, and S. C. Hicks. An integrated single-nucleus and spatial transcriptomics atlas reveals the molecular landscape of the human hippocampus. Nature Neuroscience, 28(9):1990–2004, Sept. 2025. ISSN 1546-1726. doi:10.1038/s41593-025-02022-0. URL https://www.nature.com/articles/s41593-025-02022-0.

[9] L. Huuki-Myers, A. Spangler, N. Eagles, K. D. Montgomery, S. H. Kwon, B. Guo, M. Grant-Peters, H. R. Divecha, M. Tippani, C. Sriworarat, A. B. Nguyen, P. Ravichandran, M. N. Tran, A. Seyedian, PsychENCODE consortium, T. M. Hyde, J. E. Kleinman, A. Battle, S. C. Page, M. Ryten, S. C. Hicks, K. Martinowich, L. Collado-Torres, and K. R. Maynard. Integrated single cell and unsupervised spatial transcriptomic analysis defines molecular anatomy of the human dorsolateral prefrontal cortex, Feb. 2023. URL http://biorxiv.org/lookup/doi/10.1101/2023.02.15.528722.

[10] K. R. Maynard, L. Collado-Torres, L. M. Weber, C. Uytingco, B. K. Barry, S. R. Williams, J. L. Catallini, M. N. Tran, Z. Besich, M. Tippani, J. Chew, Y. Yin, J. E. Kleinman, T. M. Hyde, N. Rao, S. C. Hicks, K. Martinowich, and A. E. Jaffe. Transcriptome-scale spatial gene expression in the human dorsolateral prefrontal cortex. Nature Neuroscience, 24(3):425–436, Mar. 2021. ISSN 1097-6256, 1546-1726. doi:10.1038/s41593-020-00787-0. URL https://www.nature.com/articles/s41593-020-00787-0.

[11] H. L. Crowell, Y. Dong, I. Billato, P. Cai, M. Emons, S. Gunz, B. Guo, M. Li, A. Mahmoud, A. Manukyan, H. Pagès, P. Panwar, S. Rao, C. J. Sargeant, L. Shepherd Kern, M. Ramos, J. Sun, M. Totty, V. J. Carey, Y. Chen, L. Collado-Torres, S. Ghazanfar, K. D. Hansen, K. Martinowich, K. R. Maynard, E. Patrick, D. Righelli, D. Risso, S. Tiberi, L. Waldron, R. Gottardo, M. D. Robinson, S. C. Hicks, and L. M. Weber. Orchestrating Spatial Transcriptomics Analysis with Bioconductor, Nov. 2025. URL http://biorxiv.org/lookup/doi/10.1101/2025.11.20.688607.

[12] P. Bankhead, M. B. Loughrey, J. A. Fernández, Y. Dombrowski, D. G. McArt, P. D. Dunne, S. Mc-Quaid, R. T. Gray, L. J. Murray, H. G. Coleman, J. A. James, M. Salto-Tellez, and P. W. Hamilton. QuPath: Open source software for digital pathology image analysis. Scientific Reports, 7(1):16878, Dec. 2017. ISSN 2045-2322. doi:10.1038/s41598-017-17204-5. URL https://www.nature.com/articles/s41598-017-17204-5.

[13] R. S. Bivand, E. Pebesma, and V. Gómez-Rubio. Applied Spatial Data Analysis with R. Springer New York, New York, NY, 2013. ISBN 9781461476177 9781461476184. doi:10.1007/978-1-4614-7618-4. URL https://link.springer.com/10.1007/978-1-4614-7618-4.

[14] D. J. McCarthy, K. R. Campbell, A. T. L. Lun, and Q. F. Wills. Scater: pre-processing, quality control, normalization and visualization of single-cell RNA-seq data in R. Bioinformatics, 33(8): 1179–1186, Apr. 2017. ISSN 1367-4803, 1367-4811. doi:10.1093/bioinformatics/btw777. URL https://academic.oup.com/bioinformatics/article/33/8/1179/2907823.

[15] R. J. Hijmans, A. Brown, and M. Barbosa. terra: Spatial Data Analysis, Mar. 2020. URL https://CRAN.R-project.org/package=terra.

